# Influence of climate forecasts, data assimilation, and uncertainty propagation on the performance of near-term phenology forecasts

**DOI:** 10.1101/2020.08.18.256057

**Authors:** Shawn D. Taylor, Ethan P. White

**Author notes:** Corresponding author: Shawn D. Taylor.

## Abstract

Evaluation of ecological forecasts is a vital step in the continuous improvement of near-term ecological forecasts. Here we performed a thorough evaluation of a near-term phenological forecast system which has been operating for several years. We evaluated point forecast accuracy and the reliability of the prediction intervals. We also tested the contribution of upstream climate forecasts on phenology forecast proficiency. We found that 9 month climate forecasts contributed little skill overall, though some species did benefit from them. The assimilation of observed winter and spring temperature provided the largest improvement of forecast skill throughout the spring. We also found that phenology forecast prediction intervals were most robust when uncertainty was propagated from climate, phenological model, and model parameters as opposed to using climate uncertainty alone. Our analysis points the way toward several potential improvements to the forecasting system, which can be re-evaluated at a future date in a continuous cycle of forecast refinement.

## INTRODUCTION

Predicting the future state of ecological systems is essential for conservation, management, stakeholder engagement, and the evaluation of ecological models. Near-term iterative forecasting, where forecasts are regularly updated based on the newest available data, can improve the accuracy of future predictions by using up-to-date information about the system and the newest forecasts for the drivers of the system (Dietze et al., 2018; White et al., 2019). The regularly updated nature of these forecasts also allows stakeholders and decision makers to use these forecasts for decision making (Dietze et al., 2018). One area of ecology with clear applications for near-term iterative forecasting is plant phenology, the timing of regularly occurring events including leaf-out, flowering, and fruiting. The timing of these events has broad influences on ecosystems and is relevant to agriculture, tourism, and wildlife management.

While near-term iterative ecological forecasting is becoming more prevalent (Van Doren and Horton, 2018; Welch et al., 2019; White et al., 2019; Thomas et al., 2020; Pearlstine et al., 2020) there has been limited work evaluating these forecasts and attempting to determine how different aspects of the iterative system influence their performance. Quantifying the proficiency of a forecast and how it is influenced by different aspects of data and modeling aids end users in forecast interpretation, allows inter-comparison of different models and methods, and can point the way toward model improvement. Three key aspects of iterative forecasts are driver forecasts (e.g., forecasts for future weather/climate), data assimilation (updating models with the newest information on drivers or ecological responses or both), and uncertainty propagation (incorporating uncertainty from each step in the forecast into the final predicted uncertainty). In the context of phenology this means a focus on forecasts and data assimilation of temperature drivers and the propagation of uncertainty from these climate drivers, the choice of phenology model, and the parameters of the model.

The use of long-term driver forecasts (greater than 25 years) based on climate scenarios is ubiquitous in ecological research, but near-term climate and weather forecasts (less than 1 year) have received less attention. The degree to which near-term weather and climate forecasts improve near-term ecological forecasts is currently unclear, and incorporating these forecasts into ecological models can be difficult due to computational challenges (Taylor and White, 2020) and substantial uncertainty in the forecasts (Dietze, 2017). This uncertainty, coupled with the potentially chaotic nature of some ecological responses (Perretti et al., 2013), may limit the effectiveness of integrating these climate and weather forecasts. Additionally, population time-series have been shown to be well predicted one time step ahead by simple models without climate drivers (Ward et al., 2014). However, many aspects of ecological systems are closely tied to weather and climate and should benefit from including these forecasts. For example, Van Doren and Horton (2018) showed a bird migration model had ample explanatory value using a 7-day weather forecast (R2=0.62), and Carrillo et al. (2018) found that a spring index model had positive skill up to a two month lead time by integrating climate forecasts. Exploring how these forecasted drivers, and their associated uncertainty, affect performance is an important step in evaluating near-term ecological forecasts (Dietze et al., 2018).

Assimilating the most up-to-date data on these climate drivers prior to forecasting is also likely to in-fluence phenology forecasts due to the influence of lagged variables. Plants rely heavily on environmental cues throughout the late winter and early spring for the timing of dormancy release (Chuine and Régnière, 2017; Piao et al., 2019). Therefore, as time passes, and more observed weather data can be assimilated into the phenological forecasts should become more accurate. Climate and weather forecasts themselves are also more accurate at shorter lead times, thus should provide more accurate phenology forecasts as dormancy release draws nearer. From these two processes, improved climate weather forecasts at shorter lead times and assimilation of lagged variables, phenological forecasts should improve as spring and the growing season progresses. However, there has been little exploration of the relative influence of data assimilation and improved weather forecasts on ecological predictions.

Integrating climate forecasts and assimilating climate data into ecological forecast systems both come with development and computational costs (Taylor and White, 2020; Thomas et al., 2020; Welch et al., 2019), and they should be rigorously tested to justify inclusion over simpler methods. Comparison against a baseline model, such as one based on the long term climatological average, allows the assessment of “forecast skill” (Jolliffe and Stephenson, 2003; Harris et al., 2018) which indicates the improvements to the model by including detailed forecasts and data on climate.

Climate forecasts also involve additional uncertainty in the resulting ecological forecasts, and under-standing this uncertainty is essential for using forecasts for decision making (Clark, 2001; Dietze, 2017; Dietze et al., 2018). Focusing on point estimates alone causes end-users to apply their own, potentially inaccurate, uncertainty and lead to less decisive decision making (Joslyn and Savelli, 2010; Savelli and Joslyn, 2013). Communicating quantifiably derived uncertainty metrics allows for the best-informed decisions and gaining trust in a forecast (Raftery, 2016). Prediction intervals which are overly narrow (too confident) or overly wide (not confident enough) can lead to flawed decision making, especially if end-users make decisions based on probabilistic values (Zhu et al., 2002). The phenology forecast system currently used in production only uses uncertainty associated with climate, but additional uncertainty from the phenological model components may be beneficial. Specifically the uncertainty associated with choice of phenological model and the model’s parameters may affect the size and proficiency of the prediction intervals (Migliavacca et al., 2012).

Here we evaluated a near-term phenological forecast system which has been online for over 2 years (Taylor and White, 2020). We show how integrating current season temperature can improve forecasts over a climatological average, and decompose the contribution from observed versus forecasted climate. The uncertainty of the forecast system was also evaluated, and partitioned into the respective sources the model, model parameters, and climate drivers. Finally we discuss how best to implement these findings and the implications to other forecast systems.

## METHODS

Detailed description of the phenology forecast system and its models are described in Taylor and White (2020). In brief, for each species and phenophase an ensemble of four phenology models was developed using data from the National Phenology Network (USA National Phenology Network, 2019) and temperature from PRISM (PRISM Climate Group, 2004). At each forecast issue date observed daily temperature from the PRISM dataset is combined with downscaled NOAA CFSv2 forecasts (Saha et al., 2014). The combined dataset is used with with each of the phenological models to produce a predicted day of year (DOY) for the onset of the specified event (ie. budburst, flowering, mature fruit, or fall colors).

### Evaluation of Point Forecasts

The current archived operational phenological forecasts hold a point value and uncertainty based only on climate. Therefore, to allow detailed evaluation of all sources of uncertainty we retroactively performed hindcasts every four days from Dec. 1 to May 30 2018. For each issue date we obtained the five most recent climate forecasts available prior to the issue date, downscaled them according to the methods described in Taylor and White (2020), and applied the phenology models to produce a phenology hindcast. We used the same four-model ensemble (Linear, Alternating, ThermalTime, Uniforc) described in Taylor and White (2020), with the exception that the ensemble is unweighted and each of the four models was fit 50 times using bootstrapping of the data. This allows us to estimate both model and parameter uncertainty.

For evaluation we obtained all USA-NPN observations from the 2018 growing seasons (Jan. 1 to June 1, 2018) and filtered them following protocols in Taylor and White (2020). Using the coordinates of each observation we extracted the hindcasts for each issue date. Thus for each observation there are hindcasts from 48 issue dates throughout the prior winter and spring. For each issue date we calculated the RMSE and coverage (see below) of all observations. We also calculated the RMSE for the four most abundant taxon (three distinct species and one species complex) observed in the spring of 2018.

### Evaluation of Uncertainty

We evaluated the uncertainty in our forecast system by calculating the coverage, which is the fraction of observations that fall within the prediction interval. For a forecast with uncertainty based on a 95% prediction interval perfect coverage is obtained when 95% of the observations fall within those intervals. A coverage below 95% signifies the forecast is too confident, and above 95% not confident enough (Harris et al., 2018). Uncertainty in our phenological forecasts can come from three sources: 1) uncertainty in forecasted climate, 2) uncertainty in selecting the true phenological model that represents the underlying mechanisms, and 3) uncertainty in the parameters fit for the selected models (Dietze, 2017). We looked at the additive contribution of uncertainty from these three sources by calculating the coverage on three different posterior distributions for each hindcast prediction at each issue date. The first represents climate uncertainty and is the standard deviation of the mean prediction for each climate ensemble member. The second is climate and model uncertainty and is the standard deviation of the mean prediction for each phenology model within each climate ensemble member. The third combines all sources (climate, model, and parameter) and is the standard deviation of the full posterior among all climate members, phenology models, and bootstrapped parameter sets (see Fig. S1).

### Evaluating Changes in Forecast Accuracy With Issue Date

Forecast accuracy is expected to decrease as the lead time (or forecast horizon) of the forecast increases (i.e., forecasts further into the future are generally less accurate; Petchey et al. (2015)). Calculating the forecast horizon is complicated for large scale phenology forecasts because the response variable itself is a date that varies across space, confounding the traditional concept of lead time. For example, given a forecast issued on Feb. 1, a prediction of leaf onset happening Feb. 15 could be a 15 day lead time. At a higher latitude the same Feb. 1 forecast may predict a leaf onset of March 2, resulting in a 30 day lead time. This is different from standard forecast horizon analyses which analyze predictions for the value of a non-time response a fixed number of days into the future. Given these complexities, we analyze forecast accuracy at different issue dates rather than lead times. This provides information on the overall accuracy of phenology forecasts made on different dates for the United States. We explored other approaches including calculating lead time as the difference between issue date and forecast date of the phenological event, but all of these solutions introduced complexities that made interpretation difficult. Future development of methods for analyzing lead time of date predictions would improve research on phenology forecasts.

### Influence of data assimilation and near-term temperature forecasts on forecast skill

To evaluate and compare the contribution to forecast skill derived from near-term temperature forecasts and data assimilation of observed temperatures, we compared three distinct approaches to generating future temperatures for the phenological forecasts (Fig. 1). The first method combines both data assimilation of observed temperature (from PRISM) up to the issue date and near-term temperature forecasts from five member climate forecast ensemble (Method 1, Fig. 1A). For each phenological model this produces five predictions (one for each climate ensemble member) which are used to produce an average prediction date along with associated climate uncertainty from the variance among the five climate ensemble members. This is the method currently implemented in the automated forecasting system. The second method uses data assimilation but replaces near-term forecasts with historical information on temperature to provide a baseline to assess the value of near-term forecasts. This is done by assimilating observed temperature up to the specified issue date and replacing the five near-term temperature forecasts from NOAA with historic observed temperature time series from each of the last 20 years. For example for an issue date of Feb. 1, 2018 the prediction uses the observed temperature from PRISM for Nov. 1, 2017 thru Jan. 31, 2018 and the future temperatures are modeled as historic temperature from Feb. 1 - Aug. 1, 2018 for each historic year from 1996-2015. For each phenological model this produces twenty predictions (one for each year from 1996-2015) at each issue date, which are used to produce a mean prediction date along with associated climate uncertainty from the variance among the 20 different years (Method 2, Fig. 1B). This method allows us to discern the contribution of observed versus forecast temperature to phenology forecast performance. The final method uses neither climate data assimilation nor near-term temperature forecasts. This is done by replacing both observed temperatures up to the issue date and forecast temperatures beyond the issue date with historic observed temperature time series from 1996-2015. For each phenological model this produces twenty predictions (one for each from 1996-2015), which are used to produce a mean prediction along with associated climate uncertainty from the variance among the twenty different historical time series (Method 3, Fig. 1C).

**Figure 1.**
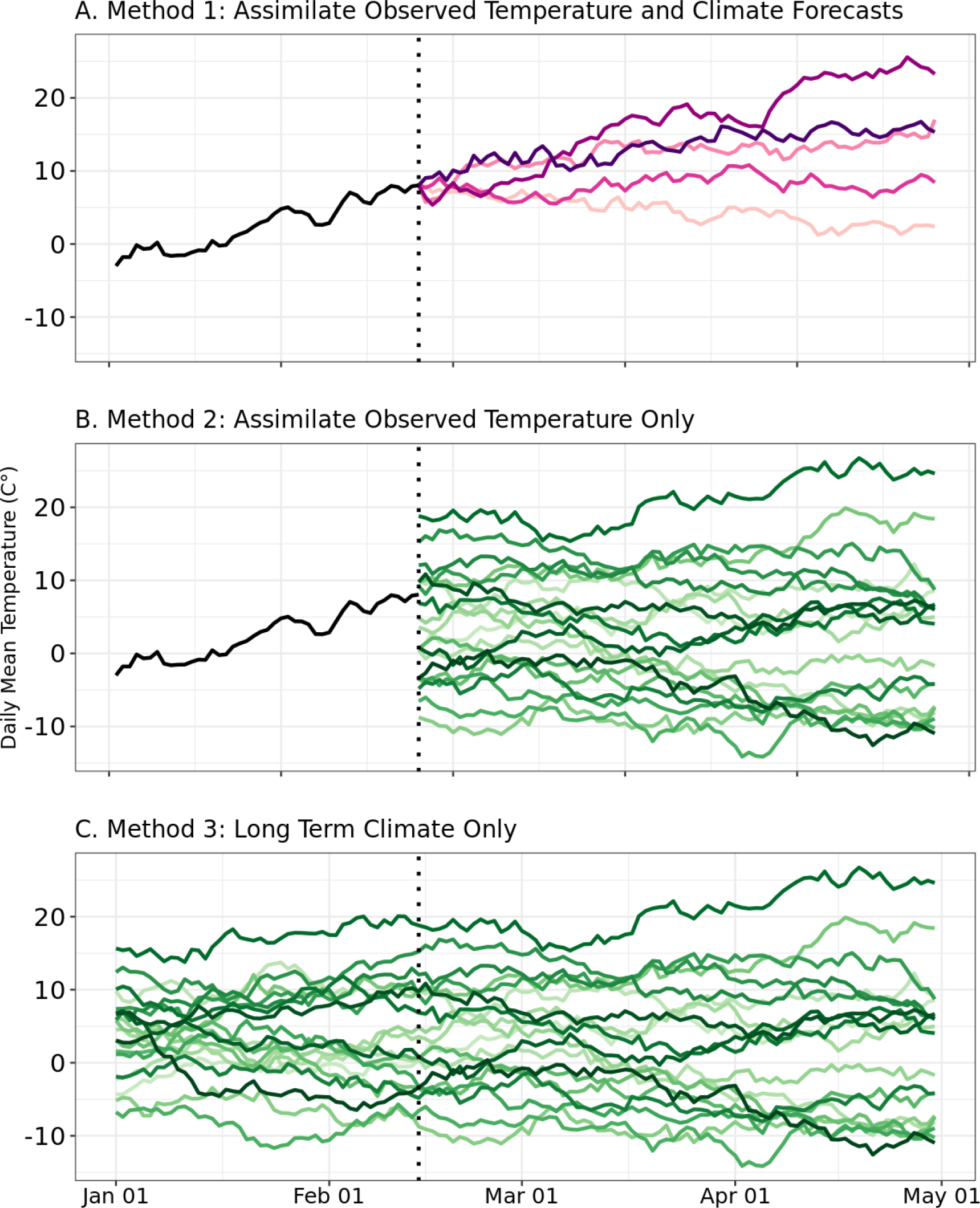
Simulated example showing the three different approaches to climate data integration for both observed (pre-issue date) and forecast (post-issue date) temperatures. The dotted vertical line indicates the issue date. Black lines indicate observed temperatures from the year of the forecast up to the issue date is used. Method 1 (A) uses 5 climate forecast members (purple lines) each integrated with observed temperature. Method 2 (B) uses 20 historic climate members for forecasts (green lines; historic data from from 1996-2015), which are each integrated with observed temperature up to the issue date. Method 3 (C) uses only data on historic climate, using 20 historical time series of observed temperature for both the observed and forecast temperatures.

Software packages used throughout the evaluation include, for the R language, ggplot2 (Wickham, 2016), prism (Hart and Bell, 2015), tidyr (Wickham and Henry, 2018), lubridate (Grolemund and Wickham, 2011). From the python language we also utilized xarray (Hoyer and Hamman, 2017), dask (Dask Development Team, 2016), scipy (Virtanen et al., 2020), numpy (Oliphant, 2006)], and pandas (McKinney, 2010). All code described is available on a GitHub repository (https://github.com/sdtaylor/phenology_forecasts). The code as well as 2018 hindcasts and observations are also permanently archived on Zenodo (https://doi.org/10.5281/zenodo.3990010).

## RESULTS

For early issue dates (predictions for spring events made in December through March), average point estimate predictions (across species and locations) made using both data assimilation of observed temperature and near-term climate forecasts (Method 1) had larger errors that those based on data assimilation of observed temperature alone (Method 2) and those using only climatology (no data assimilation or near-term forecasts, Method 3) (Fig. 2A). Climatology and data assimilation only forecasts had very similar point estimate errors during this period (Fig. 2A). All three methods underestimated uncertainty during this period, with the assimilation+forecast method generally having the worst uncertainty estimates (Fig. 2B).

**Figure 2.**
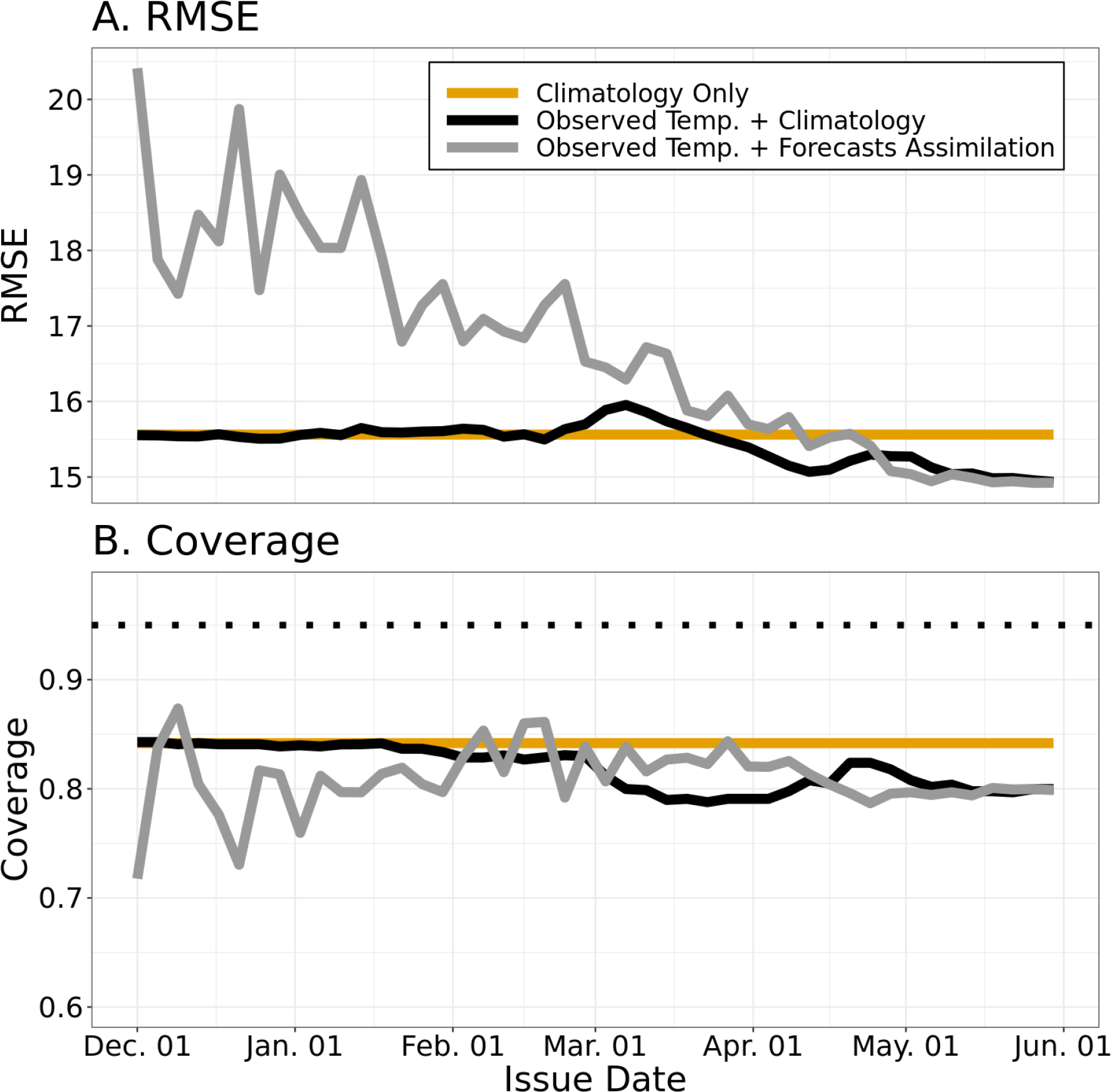
The root mean square error (RMSE) and coverage of all forecasts using the three methodologies for data assimilation. Horizontal orange lines indicate the RMSE from using the long-term climatological average. Coverage was calculated using all three sources of uncertainty (climate, model, and parameter).

Beginning around April 1, average point estimates based on assimilation-only and assimilation+forecast methods produced similar errors that were lower than the errors from climatology alone (Fig. 2A). However uncertainty estimates were better (higher coverage) for the climatology based predictions during this period (Fig. 2B). For a short period from the beginning of March to mid-April coverage from observed temperature and climatology was the worst overall.

We also explored patterns within individual species by focusing on the four taxon with the most data in the National Phenology Network dataset (Fig. 3). *Acer rubrum* showed similar patterns to the across species average. *Forsythia spp*. had similar errors for all three methods for early issue dates, the best performance by the assimilation+forecast method for intermediate issue dates, and equivalent performance by assimilation+forecast and assimilation-only methods for late issue dates (with worse predictions by the climatology method). *Prunus serotina* showed a pattern similar to *Forsythia spp*., but with the assimilation+forecast method performing slightly worse than the other method for early issue date forecasts. For *Cornus florida* the errors were highest for the assimilation+forecast method until the last few issue dates.

**Figure 3.**
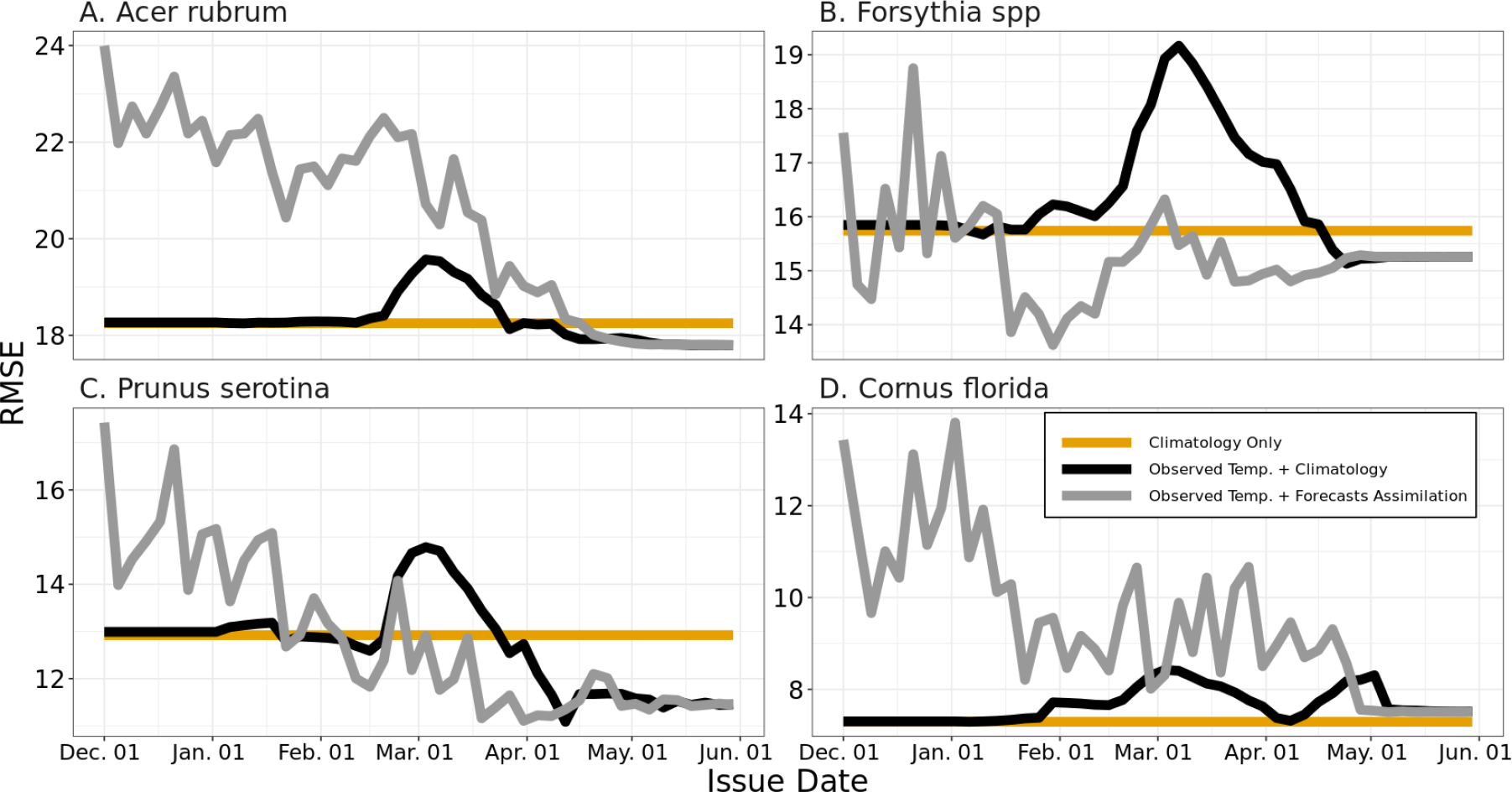
The root mean square error at different issue dates for the four taxon with the most observed data points in USA-NPN network dataset in the spring of 2018.

Across all three methods, incorporating all sources of uncertainty (climate, model, and parameter) led to the best coverage at approximately 80% of observed data points falling within the 95% prediction interval (Fig. 4). When prediction intervals were derived using only climate uncertainty, then coverage decreased to nearly 0 by the end of the forecast period, meaning that failing to account for other sources of uncertainty would have resulted in dramatically overconfident predictions. When prediction intervals were derived using only climate and model uncertainty coverage decreased slightly and was consistently worse than using all three uncertainty sources together.

**Figure 4.**
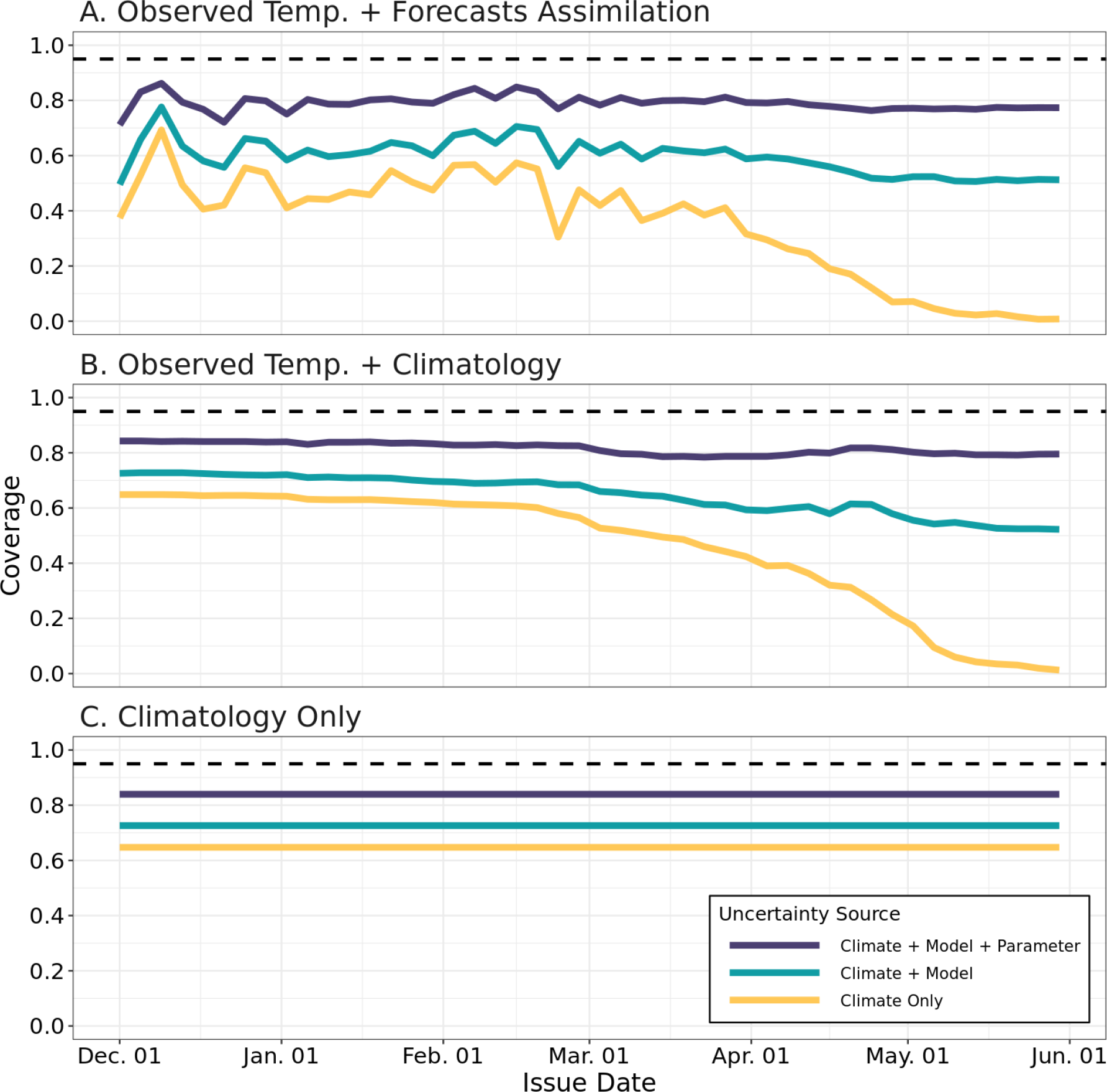
The average coverage for all forecasts. The dashed line indicates perfect coverage, 0.95, when using a 95% prediction interval. Panels indicate the three methods for climate data integration; current year observed temperature and forecast temperature (Method 1; A), current year observed temperature and climatology (Method 2; B), and climatology only (Method 3; C). Colors indicate the coverage when different sources of uncertainty are propagated.

## DISCUSSION

We evaluated a near-term species level plant phenology forecast system to explore the importance of data assimilation, near-term climate forecasts, and uncertainty propagation in the performance of the forecasts. Evaluation is an important step as it gives users a quantitative understanding of forecast performance. Our analysis showed that plant phenology forecasts for 2018 were skillful (better than the long-term average) starting in mid-April on average. Most of the skill came from assimilating observed temperature data prior to making the forecast, not from using forecasted temperature. While overall performance across all species was not improved by incorporating climate forecasts, several species did benefit from them. Uncertainty in forecasts, though slightly overconfident, was best when propagating all three sources of uncertainty (climate, model, and parameter) (Dietze et al., 2018). These results illustrate the central importance of assimilating continuously updated climate data into automated systems for making ecological forecasts and suggests that further exploration into the best ways to integrate near-term climate forecasts into phenological prediction is necessary.

While several studies have shown ecological forecasting can be aided by weather and climate forecasts (Carrillo et al., 2018; Van Doren and Horton, 2018), climate forecasts with a lead time of 1 year or less are still rarely used in ecology, likely because of the associated computational challenges (Taylor and White, 2020). Our species level analyses show some potential value for incorporating these climate forecasts, which produced increases in skill for two of the most common plants in our forecast system, *Forsythia spp*. and *Prunus serotina*, midway through the spring of 2018. These were the exception though, as overall climate forecasts provided little improvement over the assimilation of observed temperature, which provided most of the increase in skill. This is surprising since temperature plays such a fundamental role in the timing of phenological events (Chuine and Régnière, 2017; Piao et al., 2019).

The minimal gains from including temperature forecasts are due in part to the high uncertainty present in near-term temperature forecasts more than a week into the future (Dietze, 2017). Our system uses temperature forecasts obtained from the CFSv2 global climate model, which makes better predictions than climatological temperature only 20% of the time from January to June (Saha et al., 2014). This uncertainty in the climate forecasts places a hard limit on how much better the resulting phenological forecasts can perform than a climatology only method. However, disregarding climate forecasts entirely is likely not desirable, as assimilating observed temperature alone exhibited increased errors midway thru spring. This increase in error was even more pronounced at the species level. This was likely due to improbable temperature profiles from combining observed and historic temperature (Fig. 1C, Fig. S2) resulting in increasingly inaccurate predictions. In these cases the integration of climate forecasts is advantageous.

Improvements could be made to the climate forecast integration, such as improved downscaling methods or the addition of alternative global climate models. Carrillo et al.(2018) found a large increase in skill by using a post-hoc ensemble bias correction with a long training time series, and a similar bias correction could be applied to in our system. There are other abiotic drivers which are important for phenology, such as precipitation and daylength (Diez et al., 2012). Integrating other drivers from climate forecast models may not be advantageous though as they generally have low skill at the needed scales. For example precipitation has very low skill in the CFSv2 (Saha et al., 2014). One option may be utilizing teleconnections, or large-scale climatic indices. The CFSv2 has very high skill in forecasting the El Nin∽o 3.4 Index. Thus, correlative models using large scale indices, as opposed to localized process based models, may be more suitable for ecological forecasts when integrating upstream climate forecasts (Hallett et al., 2004).

With the current limitations of integrating different climate data there are several potential paths forward. One option is to focus the forecasts on the data assimilation only model (Method 2) and disregard the climate forecasts. This model performs best overall with reasonable predictions at all issue dates. However, it does yield the worst predictions for some species at intermediate issue dates (Fig. 2B). Therefore, an alternative would be to disregard current year data entirely, and use only the climatological method (Method 3). This would essentially mean a static prediction representing the long term average and variation. This simple approach provides stable predictions with relatively low RMSEs (ca. 2 weeks), and providing this information at the species level across the entire range for 78 species provides actionable information for decision making. This highlights the fact that even baseline or unskillful models can be useful if all that is needed are accurate predictions (Harris et al., 2018). However, we know that for later season forecasts data assimilation and, in some cases, temperature forecasts can improve predictions. So, an immediate solution would be an ensemble of two methods (climatology only and observed temperature assimilation plus forecasted temperature) weighted by species and time of year. The inclusion of temperature forecasts should not make phenology predictions worse on average, so the decrease in performance of this model reflects variance from the average in the evaluation year. Ensembling this model with the climatology method, and shifting the weight from the climatology method to the assimilation+forecast method as spring progresses, could provide the best aspects of all different approaches.

We tested different combinations of uncertainty from three sources: a climate ensemble, a phenological model ensemble, and variation around phenological model parameters. Using only uncertainty derived from climate is not sufficient as it leads to overly confident prediction intervals, especially when climate uncertainty is low in late spring (Fig. S3). Integrating the other two sources of uncertainty allowed for more reliable, though still overconfident, prediction intervals throughout the spring. Incorporating other sources of uncertainty, such as from an observation or process model component, are future avenues for improving the reliability of phenology forecasts (Dietze, 2017). Alternatively, methods which improve the point forecasts, thereby shifting the prediction interval, will also improve forecast uncertainty.

Evaluation of forecast performance is an important step in developing reliable and trustworthy ecological forecast infrastructure (Dietze et al., 2018). Here we showed that an automated continental scale forecast system can produce skillful forecasts starting in mid-spring on average, and earlier for some species. Though most of the skill comes from assimilating observed, as opposed to forecasted, temperature. Future improvement in the forecast system can focus on better modelling and ensemble methods or integrating large scale climate indices such as sea surface temperature. Forecast prediction intervals should include all possible sources of uncertainty to improve reliability. After changes are implemented future evaluations should confirm any improvement to forecast skill using the latest observations. This continuous cycle of open evaluation and improvement will facilitate a reliability and trustworthy plant phenology forecast system.

## ACKNOWLEDGMENTS

This research was supported by the Gordon and Betty Moore Foundation’s Data-Driven Discovery Initiative through Grant GBMF4563 to E.P. White. We thank the USA National Phenology Network and the many participants who contribute to its Nature’s Notebook program.

## REFERENCES

Carrillo, C. M., Ault, T. R., and Wilks, D. S. (2018). Spring onset predictability in the north american multimodel ensemble. Journal of Geophysical Research: Atmospheres, 123(11):5913–5926.

Chuine, I. and Régniére, J. (2017). Process-based models of phenology for plants and animals. Annual Review of Ecology, Evolution, and Systematics, 48(1):159–182.

Clark, J. S. (2001). Ecological forecasts: An emerging imperative. Science, 293(5530):657–660.

Dask Development Team (2016). Dask: Library for dynamic task scheduling.

Dietze, M. C. (2017). Prediction in ecology: a first-principles framework. Ecological Applications, 27(7):2048–2060.

Dietze, M. C., Fox, A., Beck-Johnson, L. M., Betancourt, J. L., Hooten, M. B., Jarnevich, C. S., Keitt, T. H., Kenney, M. A., Laney, C. M., Larsen, L. G., Loescher, H. W., Lunch, C. K., Pijanowski, B. C., Randerson, J. T., Read, E. K., Tredennick, A. T., Vargas, R., Weathers, K. C., and White, E. P. (2018). Iterative near-term ecological forecasting: Needs, opportunities, and challenges. Proceedings of the National Academy of Sciences, 115(7):1424–1432.

Diez, J. M., Ibáñez, I., Miller-Rushing, A. J., Mazer, S. J., Crimmins, T. M., Crimmins, M. A., Bertelsen, C. D., and Inouye, D. W. (2012). Forecasting phenology: from species variability to community patterns. Ecology Letters, 15(6):545–553.

Grolemund, G. and Wickham, H. (2011). Dates and times made easy with {lubridate}. Journal of Statistical Software, 40(3):1–25.

Hallett, T. B., Coulson, T., Pilkington, J. G., Clutton-Brock, T. H., Pemberton, J. M., and Grenfell, B. T. (2004). Why large-scale climate indices seem to predict ecological processes better than local weather. Nature, 430(6995):71–75.

Harris, D. J., Taylor, S. D., and White, E. P. (2018). Forecasting biodiversity in breeding birds using best practices. PeerJ, 6:e4278.

Hart, E. M. and Bell, K. (2015). prism: Download data from the oregon prism project. http://github.com/ropensci/prism.

Hoyer, S. and Hamman, J. J. (2017). xarray: N-d labeled arrays and datasets in python. Journal of Open Research Software, 5.

Jolliffe, I. T. and Stephenson, D. B., editors (2003). Forecast verification: a practitioner’s guide in atmospheric science. John Wiley and Sons, Ltd.

Joslyn, S. and Savelli, S. (2010). Communicating forecast uncertainty: public perception of weather forecast uncertainty. Meteorological Applications, 17(2):180–195.

McKinney, W. (2010). Data structures for statistical computing in python. In Proceedings of the 9th Python in Science Conference, pages 51–56, Austin, Texas, USA. SciPy.

Migliavacca, M., Sonnentag, O., Keenan, T. F., Cescatti, A., O’Keefe, J., and Richardson, A. D. (2012). On the uncertainty of phenological responses to climate change, and implications for a terrestrial biosphere model. Biogeosciences, 9(6):2063–2083.

Oliphant, T. (2006). A guide to NumPy. Trelgol Publishing, Provo, UT.

Pearlstine, L. G., Beerens, J. M., Reynolds, G., Haider, S. M., McKelvy, M., Suir, K., Romañach, S. S., and Nestler, J. H. (2020). Near-term spatial hydrologic forecasting in everglades, usa for landscape planning and ecological forecasting. Environmental Modelling and Software, page 104783.

Perretti, C. T., Munch, S. B., and Sugihara, G. (2013). Model-free forecasting outperforms the correct mechanistic model for simulated and experimental data. Proceedings of the National Academy of Sciences, 110(13):5253–5257.

Petchey, O. L., Pontarp, M., Massie, T. M., Kéfi, S., Ozgul, A., Weilenmann, M., Palamara, G. M., Altermatt, F., Matthews, B., Levine, J. M., Childs, D. Z., McGill, B. J., Schaepman, M. E., Schmid, B., Spaak, P., Beckerman, A. P., Pennekamp, F., and Pearse, I. S. (2015). The ecological forecast horizon, and examples of its uses and determinants. Ecology Letters, 18(7):597–611.

Piao, S., Liu, Q., Chen, A., Janssens, I. A., Fu, Y., Dai, J., Liu, L., Lian, X., Shen, M., and Zhu, X. (2019). Plant phenology and global climate change: Current progresses and challenges. Global Change Biology, 25(6):1922–1940.

PRISM Climate Group (2004). Oregon state university.

Raftery, A. E. (2016). Use and communication of probabilistic forecasts. Statistical Analysis and Data Mining: The ASA Data Science Journal, 9(6):397–410.

Saha, S., Moorthi, S., Wu, X., Wang, J., Nadiga, S., Tripp, P., Behringer, D., Hou, Y.-T., Chuang, H.-y., Iredell, M., Ek, M., Meng, J., Yang, R., Mendez, M. P., van den Dool, H., Zhang, Q., Wang, W., Chen, M., and Becker, E. (2014). The ncep climate forecast system version 2. Journal of Climate, 27(6):2185–2208.

Savelli, S. and Joslyn, S. (2013). The advantages of predictive interval forecasts for non-expert users and the impact of visualizations. Applied Cognitive Psychology, 27(4):527–541.

Taylor, S. D. and White, E. P. (2020). Automated data-intensive forecasting of plant phenology throughout the united states. Ecological Applications, 30(1).

Thomas, R. Q., Figueiredo, R. J., Daneshmand, V., Bookout, B. J., Puckett, L. K., and Carey, C. C. (2020). A near-term iterative forecasting system successfully predicts reservoir hydrodynamics and partitions uncertainty in real time. bioRxiv.

USA National Phenology Network (2019). Plant and animal phenology data. data type: Individual phenometrics. 01/01/2018-12/31/2018 for region: 49.9375, -66.4791667 (ur); 24.0625, -125.0208333 (ll).

USA-NPN, Tucson, Arizona, USA. Data set accessed 05/09/2019 at http://doi.org/10.5066/F78S4N1V.

Van Doren, B. M. and Horton, K. G. (2018). A continental system for forecasting bird migration. Science, 361(6407):1115–1118.

Virtanen, P., Gommers, R., Oliphant, T. E., Haberland, M., Reddy, T., Cournapeau, D., Burovski, E., Peterson, P., Weckesser, W., Bright, J., van der Walt, S. J., Brett, M., Wilson, J., Millman, K. J., Mayorov, N., Nelson, A. R. J., Jones, E., Kern, R., Larson, E., Carey, C. J., Polat, I., Feng, Y., Moore, E. W., VanderPlas, J., Laxalde, D., Perktold, J., Cimrman, R., Henriksen, I., Quintero, E. A., Harris, C. R., Archibald, A. M., Ribeiro, A. H., Pedregosa, F., and van Mulbregt, P. (2020). SciPy 1.0: fundamental algorithms for scientific computing in Python. Nature Methods, 17(3):261–272.

Ward, E. J., Holmes, E. E., Thorson, J. T., and Collen, B. (2014). Complexity is costly: a meta-analysis of parametric and non-parametric methods for short-term population forecasting. Oikos, 123(6):652–661.

Welch, H., Hazen, E. L., Bograd, S. J., Jacox, M. G., Brodie, S., Robinson, D., Scales, K. L., Dewitt, L., and Lewison, R. (2019). Practical considerations for operationalizing dynamic management tools. Journal of Applied Ecology, 56(2):459–469.

White, E. P., Yenni, G. M., Taylor, S. D., Christensen, E. M., Bledsoe, E. K., Simonis, J. L., and Ernest, S. K. M. (2019). Developing an automated iterative near-term forecasting system for an ecological study. Methods in Ecology and Evolution, 10(3):332–344.

Wickham, H. (2016). ggplot2: Elegant Graphics for Data Analysis. Springer-Verlag New York.

Wickham, H. and Henry, L. (2018). tidyr: Easily tidy data with ‘spread()’ and ‘gather()’ functions.

Zhu, Y., Toth, Z., Wobus, R., Richardson, D., and Mylne, K. (2002). The economic value of ensemble-based weather forecasts. Bulletin of the American Meteorological Society, 83(1):73–83.

