## Supplement for "Influence of climate forecasts, data assimilation, and uncertainty propagation on the performance of near-term phenology forecasts"

Supplementary images S1 - S3

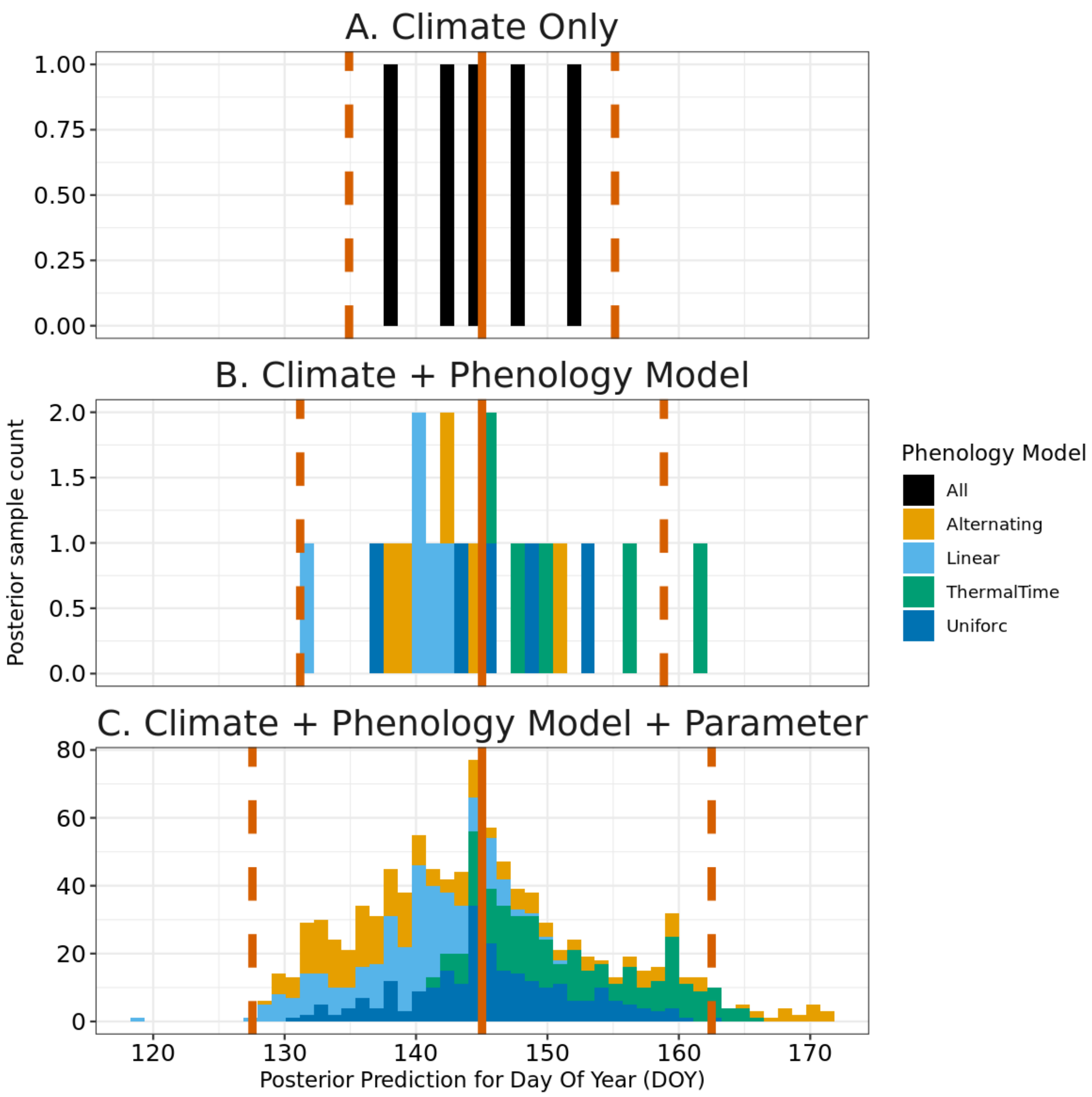


Figure S1: The posterior distribution for a single forecast. Panels represent the posterior using only uncertainty from climate (A), from climate and phenology models (B), and from all sources (C). While the mean prediction (solid red line) stays the same, the posterior 95% prediction interval (dashed red lines) is widest when uncertainty from all sources is propagated.

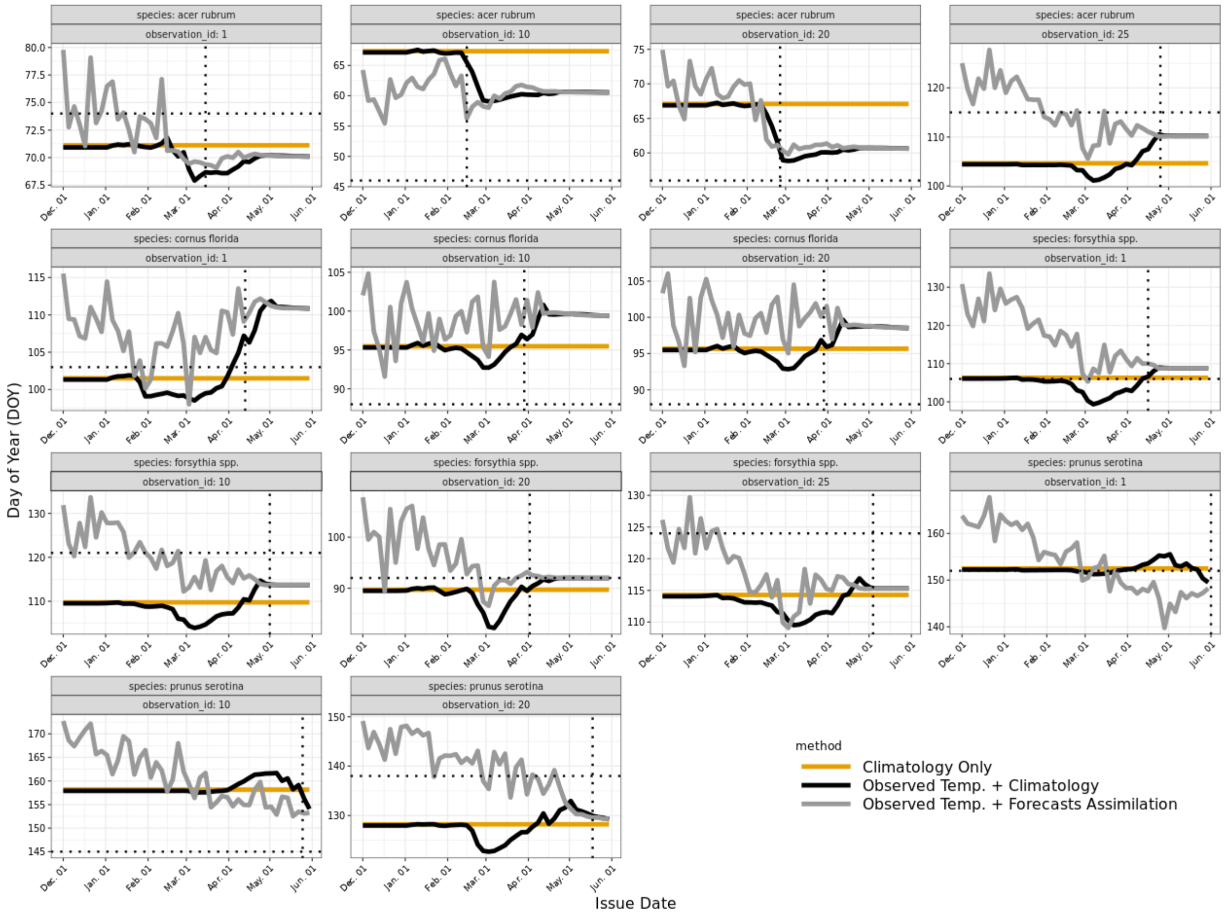


Figure S2: Actual predictions from the hindcasts of 14 budburst observations from 2018. Colors indicate the three different methods used in the analysis. Dotted lines indicate the observed budburst date on both axes.

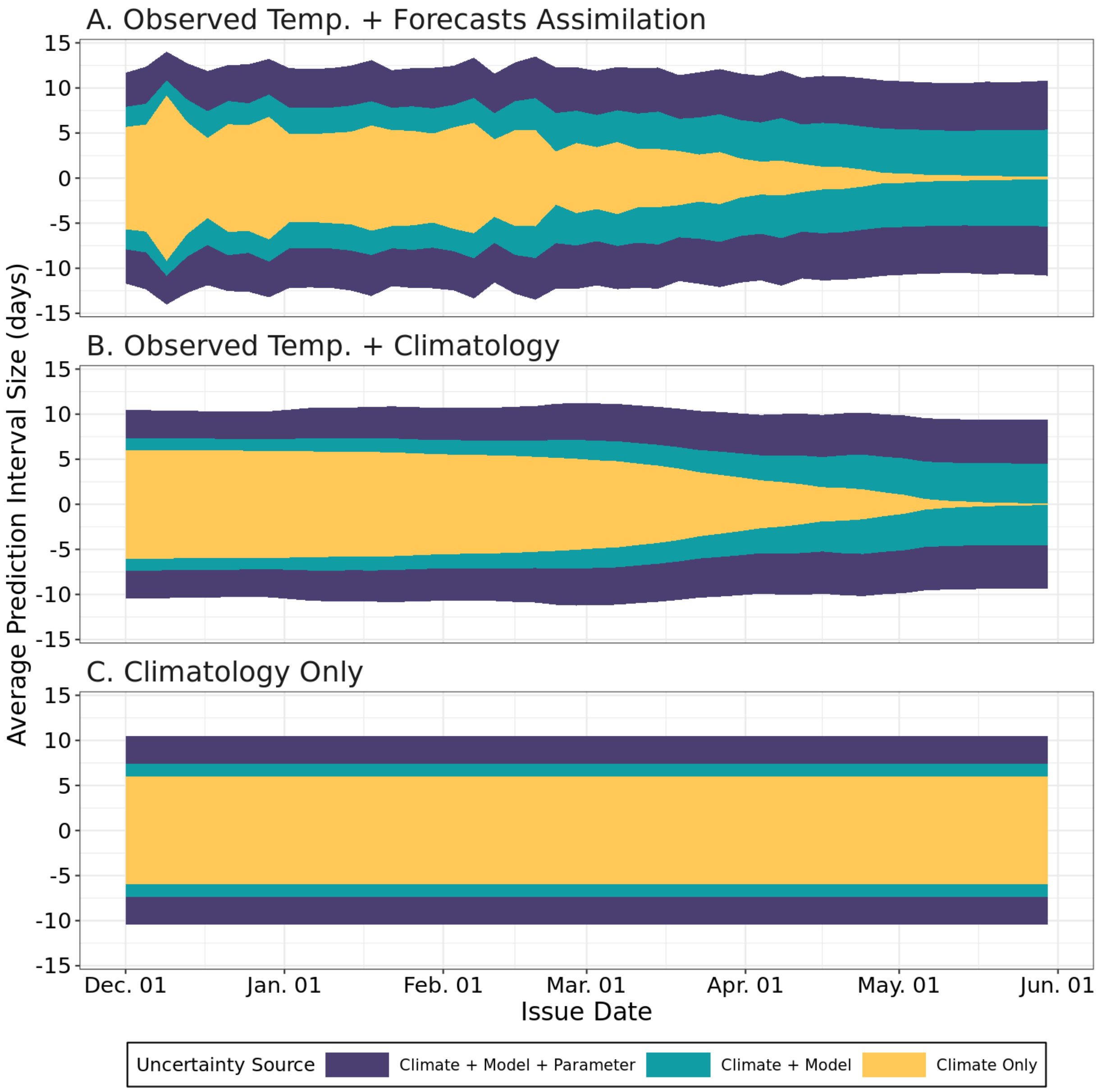


Figure S3: The average contribution to uncertainty from the different forecast components. The panels represent the three different methodologies in assimilating observed, forecast, and historical temperature. Colors indicate different attributes from the three sources of uncertainty. The y-axis indicates the average 95% confidence interval of all forecasts at each issue date from Dec. 1, 2017 to June 1, 2018.
